# The lipid raft-associated protein stomatin is required for accumulation of dectin-1 in the phagosomal membrane and for full activity of macrophages against *Aspergillus fumigatus*

**DOI:** 10.1101/2022.02.25.481948

**Authors:** Marie Goldmann, Franziska Schmidt, Zoltán Cseresnyés, Thomas Orasch, Susanne Jahreis, Susann Hartung, Marc Thilo Figge, Marie von Lilienfeld-Toal, Thorsten Heinekamp, Axel A. Brakhage

**Author notes:** These authors contributed equally to the manuscript and were listed alphabetically (Marie Goldmann, Franziska Schmidt).

## Abstract

Alveolar macrophages are among the first cells to come into contact with inhaled fungal conidia of the human pathogenic fungus *Aspergillus fumigatus*. In lung alveoli, they contribute to phagocytosis and elimination of conidia. As a counter defense, conidia contain a grey-green pigment allowing them to survive in phagosomes of macrophages for some time. Previously, we showed that this conidial pigment interferes with the formation of flotillin-dependent lipid-rafts in the phagosomal membrane thereby preventing the formation of functional phagolysosomes. In this study, the role of the lipid raft-associated protein stomatin in macrophages during antifungal defense was investigated. To determine the function of this integral membrane protein, a stomatin-deficient macrophage cell line was generated by CRISPR/Cas9 gene editing. Immunofluorescence and flow cytometry revealed that stomatin contributes to the phagocytosis of conidia and is important for recruitment of both the β-glucan receptor dectin-1 and the vATPase to the membrane of phagosomes. In the stomatin knockout cell line, fusion of phagosomes and lysosomes was reduced when infected with pigmentless *pksP* conidia. Thus, our data suggest that stomatin is involved in maturation of phagosomes *via* fostering fusion of phagosomes with lysosomes.

## Introduction

Due to advances in modern medicine, the number of immunocompromised patients has increased in recent decades. In this patient cohort a frequently diagnosed fungal infection is caused by *Aspergillus fumigatus*, a ubiquitous, saprophytic mold. The fungus produces a myriad of conidia, which are released into the atmosphere. With a diameter of 2 to 3 μm, conidia are inhaled by humans and can easily reach the lung alveoli (Tischler and Hohl, 2019, Brakhage and Langfelder, 2002). In healthy individuals, these conidia are apparently effectively cleared by the innate immune system. However, in patients with weakened immunity conidia can escape the host’s defense, and outgrowing hyphae can invade pulmonary tissue to cause invasive aspergillosis (Lee and Sheppard, 2016). The characterization of the pathogen’s defense strategies against human immune responses and the identification of relevant host factors against *A. fumigatus* are of considerable interest.

In lung alveoli, resident alveolar macrophages are part of the initial defense against inhaled conidia. The central role of macrophages is the internalization and degradation of microbial pathogens (Ibrahim-Granet et al., 2003, Aderem and Underhill, 1999). Reduced numbers or impaired activity of phagocytes, *e.g.,* due to leukemia or after stem cell transplantation, increase susceptibility of patients towards invasive aspergillosis (Brakhage, 2005, Latge, 1999, Heinekamp et al., 2015),

Macrophages recognize conidia *via* pathogen recognition receptors (PRRs) and engulf them by forming a phagocytic cup. Within the macrophage, the conidium-containing phagosome fuses with lysosomes to form a phagolysosome (Philippe et al., 2003, Heinekamp et al., 2015). This fusion process leads to an acidic pH inside the phagolysosome, which activates hydrolytic enzymes to eliminate the pathogen (Claus et al., 1998, Garin et al., 2001). The conidial surface pigment dihydroxynaphthalene (DHN) melanin plays a crucial role in host pathogen interaction. It interferes with the phagocytic uptake and inhibits phagosomal acidification (Jahn et al., 2002, Thywissen et al., 2011). In line, conidia devoid of the polyketide synthase PksP executing the first step in DHN-melanin biosynthesis (Jahn et al., 1997), are less resistant against the immune response. Melanin-free conidia show faster phagocytosis rates, increased phagolysosomal fusion events and attenuated virulence in murine infection models (Jahn et al., 1997, Jahn et al., 2002, Langfelder et al., 1998, Heinekamp et al., 2012). We previously identified the mechanism how DHN melanin interferes with the phagosome maturation: Depending on flotillins, melanin inhibits the formation of lipid-raft microdomains in the phagosomal membrane, that are required for maturation of functional phagolysosomes including assembly of NADPH oxidase and vATPase complexes (Schmidt et al., 2020).

Lipid-raft microdomains are small, heterogeneous and highly dynamic structures within cell membranes (Simons and Ikonen, 1997). These sphingolipid- and cholesterol-rich microdomains in membranes preferentially contain glycosyl phosphatidylinositol (GPI)- anchored proteins and integral proteins like flotillin-1, flotillin-2 and stomatin (Salzer and Prohaska, 2001). In general, lipid rafts play important roles in response to intracellular or extracellular stimuli (Harder et al., 1998, Biswas et al., 2008). However, the precise roles of integral proteins for formation and dynamics of lipid raft microdomains awaits further analyses. In this regard, only limited information on stomatin is available in particular on its role for defense against pathogens.

Stomatin is conserved from archaea to mammals (Lapatsina et al., 2012). Consistently with other proteins involved in membrane organization, stomatin proteins form higher order oligomers and are found enriched in lipid-raft microdomains (Snyers et al., 1999, Snyers et al., 1998). Stomatin-like and related proteins are characterized by an SPFH (stomatin, prohibitin, flotillin, HfIK/C)-domain, which is crucial for the association with the membrane scaffold and other membrane proteins (Morrow and Parton, 2005, Bauer and Pelkmans, 2006). As a major component of lipid rafts, stomatin localizes to membranes (Lee et al., 2017, Brand et al., 2012, Snyers et al., 1999, Umlauf et al., 2006, Salzer and Prohaska, 2001). To shed light on this central protein, here we characterized in detail the basic functions of stomatin in macrophages with an emphasis on its role for recognition of and defense against *A. fumigatus*.

## Materials and Methods

### Cell lines

The murine macrophage cell line RAW264.7 (ATCC TIB-71) was cultivated in DMEM (Gibco) supplemented with 10 % (v/v) fetal bovine serum (FBS) (GE Healthcare Life Sciences), 1 % (w/v) ultraglutamine (Gibco) and 27.5 μg/ml gentamicin sulfate (Gibco) at 37 °C, 5 % (v/v) CO_2_ in a humidified chamber. When required, the medium was further supplemented with puromycin (7 μg/ml for selection of clones or 6 μg/ml for maintenance of cell lines; Invivogen) and/or hygromycin (100 μg/ml for selection of clones or 70 μg/ml for maintenance of clones; Invivogen). The human kidney epithelial cell line Lenti-X™ 293T (Takara) was cultivated in DMEM supplemented with 10 % (v/v) FBS, 1 % (w/v) ultraglutamine, 1 % (v/v) sodium bicarbonate (Gibco), 1 % (v/v) penicillin/streptomycin (Gibco).

### Cultivation, staining and fixation of *A. fumigatus*

The *A. fumigatus* strains ATCC46645 (wild type) and the corresponding pigmentless *pksP* mutant (Jahn et al., 1997) were cultivated on *Aspergillus* minimal medium (AMM) agar plates (Pontecorvo et al., 1953). *A. fumigatus* was grown for 5 days at 37 °C and conidia were harvested and labeled fluorescein isothiocyanate (FITC; Sigma-Aldrich) or calcofluor white (CFW; Sigma-Aldrich) as previously described (Thywissen et al., 2011).

### Infection experiments

Macrophages were seeded in 24-well plates with cover slips at a density of 3 x 10^5^ cells/well and inspected microscopically. For further analysis of isolated phagosomes 4 x 10^6^ cells/well were seeded in 4-well plates. The cells were infected with conidia at a multiplicity of infection (MOI) of 2 or 5, as indicated. To synchronize infection, cells were incubated at 4 °C for 30 min or, alternatively, by centrifugation for 5 min at 200 x g. To start infection, cells were transferred to a humidified chamber and incubated at 37 °C for 30 min or 2 h, as indicated..

### Generation of a RAW264.7 stomatin knockout cell line

To perform gene editing in RAW264.7 macrophages, the Lenti-X CRISPR/Cas9-System (Takara) was used. First, RAW264.7 cells with stable production of Cas9 were generated according to the manufacturer’s protocol. Lenti-X 293T cells were transfected with vector pLVX-puro-Cas9 to produce lentivirus encoding the Cas9 gene. Lentiviral titer was determined by using the Lenti-X p24 Rapid Titer Kit (Takara). Then, RAW264.7 macrophages were transduced with Cas9-lentivirus particles at an MOI of 5 and monoclonal cells were obtained by selection with puromycin (7 μg/ml). To target the stomatin (*Stom*) gene, single-guided RNAs (sgRNAs) were designed with the online tool ChopChop (https://chopchop.cbu.uib.no/). Oligonucleotides CACCGAGAGTCATCATCTTTAGAC and AAACGTCTAAAGATGATGACTCTC corresponding to the target sgRNA were annealed and cloned into vector pLVX-hyg-sgRNA. Lenti-X™ 293T cells were transfected with the resulting vector pLVX-hyg-Stom-sgRNA. As a result, they produced lentivirus particles encoding *Stom*-sgRNAs. Then, RAW264.7 macrophages with stable production of Cas9 were transduced with isolated *Stom*-sgRNA-lentivirus particles. Cell clones were obtained by selection of cells with hygromycin (100 μg/μl). After several passages of cell clones, stable clones were screened for gene editing event resulting in the lack of stomatin production. A knockout of stomatin was verified by Western blotting. On the DNA level, deletion of a single nucleotide in the stomatin gene was verified by DNA sequence analysis. For this purpose, a DNA fragment comprising the sgRNA target region was amplified by PCR using primers AATAGAGCAAACAACAGGAGGC and GAGTACTGACCTGGTCCTTTGG and the Phusion Flash High-Fidelity PCR Master Mix (Thermo Scientific).

### Isolation of murine bone marrow-derived macrophages (BMDMs)

C57BL/6J mice were supplied by Charles River (Sulzfeld, Germany). Bone marrow cells from C57BL/6J and Flot1^-/-^ mice (Bitsikas et al., 2014) were isolated from femurs and tibias of 12-16 weeks-old mice following a previously described method (Zhang et al., 2008). Isolated bone marrow was treated with ACK lysis buffer (Gibco) to lyse red blood cells. BMDMs were differentiated as described previously (Schmidt et al., 2020).

### Cell lysate preparation and isolation of *Aspergillus*-containing phagosomes

To obtain protein extracts, cells were lysed with radioimmune precipitation assay (RIPA) buffer (Sigma-Aldrich). For this purpose, 1 x 10^6^ cells were sedimented by centrifugation, resuspended in 100 μl RIPA lysis buffer and centrifuged for 15 min at 16,000 x *g* and 4 °C. The supernatant contains cellular proteins.

The isolation of conidia-containing phagosomes from RAW264.7 macrophages or primary macrophages was performed according to a previously established protocol (Goldmann et al., 2021).

### SDS-PAGE and Western blot analysis

Protein concentrations were determined by Bradford assay as described (Kruger, 1994). Protein extracts of an equal amount were separated by SDS-PAGE (NuPAGE 4-12 % (w/v) Bis-Tris protein gels, Invitrogen) and transferred to a PVDF membrane (iBlot2 PVDF Mini Stacks, Invitrogen). The membrane was blocked with 5 % (w/v) milk powder at room temperature for 1 h and incubated overnight at 4 °C with the primary antibody. As primary antibodies, mouse anti-Cas9 antibody (MA1-201; Invitrogen), rabbit anti-Stomatin (ab166623, Abcam) and rabbit anti-vimentin as loading control (5741, Cell Signaling Technology) were used with a dilution of 1:500 (Cas9) or 1:2000 (stomatin, vimentin). After incubation with horseradish peroxidase (HRP)-conjugated IgG antibodies, goat anti-rabbit IgG-HRP (ab6721, Abcam) or anti-mouse IgG-HRP (7076, Cell Signaling Technology), both 1:2000 diluted, were added for 1 h at room temperature. Detection was performed using the 1-Step™ Ultra TMB-Blotting Solution (Thermo Scientific).

### Quantitation of the phagocytosis ratio, acidification of phagosomes and recruitment of lipid-raft microdomains

To analyze the phagocytosis ratio, macrophages were infected with FITC-labeled wild-type or *pksP* conidia with an MOI of 5. To synchronize phagocytosis, samples were immediately incubated at 4 °C for 30 min. Then, phagocytosis was started by incubation in a CO_2_ incubator at 37 °C for 30 min. The process was stopped by washing with 1 ml ice cold PBS. Non-phagocytosed conidia were counterstained with 0.25 mg/ml CFW for 15 min. Cells were washed twice with PBS and afterwards fixed for 15 min with 3.7 % (v/v) formaldehyde/PBS. Following a final washing step, coverslips were transferred to glass slides and imaged by an LSM 780 confocal microscope (Zeiss). In each biological replicate circa two hundred conidia were considered and the quantity of phagocytosed conidia was compared to the number of non-phagocytosed conidia (Schmidt et al., 2020).

For measuring acidification of phagosomes prior to infection with conidia, macrophages were prestained with 50 nM LysoTracker Red DND-99 (Life Technologies) for 1 h. Then, the medium was exchanged by fresh medium with LysoTracker Red DND-99. Macrophages were infected with CFW-labeled wild-type or *pksP* conidia with an MOI of 2 and phagocytosis was synchronized by centrifugation. After 2 h of co-incubation, cells were fixed with 3.7 % (v/v) formaldehyde/PBS as described before and imaged by a LSM 780 (Zeiss). For the evaluation of the phagosomal acidification, one hundred conidia-containing phagosomes were analyzed and a strong red signal around the conidia indicated a decrease in pH. For further characterization of the phagosomal acidification, a second dye was applied. Macrophages were preloaded with 1 μM LysoSensor Yellow/Blue DND-160 for 30 min. Afterwards the medium was exchanged by fresh medium with LysoSensor Yellow/Blue DND-160 and cells were infected with conidia (MOI of 2) as described before. After fixation, the samples were microscopically analyzed for acidification of the phagosomes. This was indicated by a shift of the emission spectrum from blue (neutral pH) to yellow (acidic pH).

To visualize GM1 (monosialotetrahexosylganglioside) in phagosomal membranes, macrophages were incubated with 1.5 μM Alexa Fluor 647-conjugated Cholera Toxin B (CTB; Life Technologies) for 1 h and afterwards infected with FITC-labeled *A. fumigatus* conidia with an MOI of 2. Phagocytosis was allowed to proceed for 2 h after synchronization by centrifugation. For microscopic analysis, cells were fixed with 3.7 % (v/v) formaldehyde/PBS as described before. One hundred conidia-containing phagosomes were evaluated for GM1 recruitment, visualized by an accumulation of the CTB-Alexa Fluor 647 dye in the phagosomal membrane. All values represent mean ±SD of three biological replicates.

### Immunofluorescence and microscopy

Cells were fixed with 3.7 % (v/v) formaldehyde/PBS for 15 min and then permeabilized with 0.1 % (v/v) Triton X-100/PBS for 10 min. Isolated phagosomes were not permeabilized. Macrophages and isolated phagosomes were blocked with 5 % (v/v) normal goat serum (Thermo Fisher Scientific), 2 % (w/v) BSA, 0.3 M glycine/PBS for 30 min before overnight incubation at 4 °C with the primary antibody in 2 % (w/v) BSA/PBS. As primary antibodies, rabbit anti-Stomatin (ab166623, Abcam), rabbit anti-ATP6V1B2 (ab73404, Abcam) or goat anti-dectin-1/CLEC7 (AF1756, R&D Systems) were used with a concentration of 1:500 (Stomatin, vATPase) or 1:100 (dectin-1). The incubation with secondary antibodies, goat anti-rabbit IgG DyLight 633 (35562, Thermo Fisher Scientific) or donkey anti-goat Alexa Fluor 488 (A11055, Life Technologies) was carried out at room temperature for 1 h with concentrations of 1:500 or 1:200. Samples were finally visualized employing a Zeiss LSM 780 confocal microscope and ZEN software (Zeiss).

### Measuring phagosomal maturation

To monitor the phagosomal maturation, macrophages were seeded with a density of 3 x 10^5^ cells/well on coverslips and stained overnight with 60 μg/ml tetramethylrhodamine dextran (3000 MW, Invitrogen). At the next day, the medium was exchanged to standard cultivation medium and cells were infected with FITC-labeled *A. fumigatus* conidia with an MOI of 2. After synchronization of phagocytosis by centrifugation and incubation for 2 h at 37 °C in a CO_2_ incubator, fusion of phagosomes and lysosomes was evaluated by confocal microscopy. Fusion was detected by an increased accumulation of tetramethylrhodamine dye at the conidia-containing phagosome. For quantification, the mean fluorescence intensity of one hundred phagosomes was calculated with the Zen software of the LSM 780 microscope (Zeiss).

### Flow cytometry

To analyze dectin-1 receptor expression on the surface of macrophages by flow cytometry, 10^6^ cells were pelleted in a 1.5 ml tube by centrifugation for 5 min at 300 x *g* and 4°C. The pellet was washed once with 1 % (w/v) BSA in PBS + 0.2 mM EDTA (FACS buffer) and subsequently resuspended in 100 μl FACS buffer containing 10 % (v/v) FcR blocking reagent (Miltenyi Biotec) and 1 % (v/v) Viobility 405/520 Fixable Dye (Miltenyi Biotec). The cell suspension was incubated for 10 min on ice following addition of 0.5 μg of either the anti-dectin-1 antibody (PE anti-mouse CD369 (dectin-1/CLEC7A) Antibody; BioLegend) or the respective isotype control (PE Rat IgG1, κ Isotype Ctrl Antibody; BioLegend) and further incubation on ice for 30 min. After this incubation time, the cells were washed 3 times by centrifugation (at 4 °C and 300 x *g* for 5 min) and resuspension in 100 μl FACS buffer. After the last washing step, the cells were resuspended in 200 μl FACS buffer and measured by flow cytometry (BD LSR Fortessa, BD Biosciences). Data were analyzed by FlowJo software (BD Biosciences).

### Image analysis

Isolated phagosomes were imaged in transmitted light (TL) and fluorescence mode of confocal microscopy to reveal the localization of the phagosomes and their vATPase level. Images were analyzed with a custom designed algorithm written in the graphical programming language JIPipe (jipipe.org). The targeted measurements consider the intensity of the vATPase fluorescence signal, detected within the phagosomes. The phagosomes were identified from the TL images, which provided a label-free way to reliably segment all phagosomes independently of their vATPase fluorescence signal. The fluorescence signal specific for vATPase was not suitable for such segmentation, because the production of the proteins of the complex varied according to the experimental conditions, thus could not be algorithmically identified. At the same time, the TL images provided a suitable platform for the segmentation process. On the other hand, TL images did not allow the segmentation of the isolated phagosomes based on pixel intensity, because the grey value of the image pixels within the phagosome areas varied according to laser intensity and relative light phase. Thus, we utilized Cellpose (Stringer et al., 2021), a Deep Learning (DL)-based generalized image segmentation framework that was originally trained on a very large number of cells and cell-like objects. Our analysis workflow was written entirely in JIPipe (Supplementary Figure 1), using specific nodes dedicated to training (Supplementary Figure 2, Supplementary Figure 5, Supplementary Figure 6) and applying Cellpose tools (Supplementary Figure 3, Supplementary Figure 7, Supplementary Figure 8) directly within the workflow. Cellpose can be applied either by using its default pre-trained model, or by training the underlying DL network either from scratch or *via* transfer learning. We combined the application of the default model and the transfer learning-based modified model. The application of the latter technique was necessary because the default model sometimes failed either by missing the darker toned phagosomes (Supplementary Figure 4A-B) or by producing false positive predictions (Supplementary Figure 4C-D), whereas the custom-trained model performed very well at the problematic images (Supplementary Figure 4E-F).

First, all images were tested with the default Cellpose model (Supplementary Figure 3, Supplementary Figure 7). After examining the outcome, 1031 isolated phagosomes were manually outlined from a subset of the images where the default model failed, in order to provide a training database for Cellpose transfer learning (Supplementary Figure 2, Supplementary Figure 5, Supplementary Figure 6). Finally, both the default and the trained models were applied in parallel (Supplementary Figure 3, Supplementary Figure 7) and the best fitting model was selected for each image (Supplementary Figure 3, Supplementary Figure 4, Supplementary Figure 8). The comparison of the two models indicated the success of this combined approach (Supplementary Figure 4).

### Quantification and statistical analysis

If not stated otherwise, at least 100 events per sample of three biological replicates were analyzed. Data are presented as mean ± SD. *P* values were calculated by a two-tailed Student’s *t* test (*, *p* < 0.05)...

## Results

### Phagosomal membranes of RAW264.7 macrophages and BMDMs contain stomatin

In general, lipid rafts are membrane microdomains rich in sphingolipids and cholesterol. They not only containing GPI-anchored proteins but also integral membrane proteins like flotillins and stomatin (Salzer and Prohaska, 2001). Since the function of lipid-raft microdomains for phagosome biogenesis is not yet fully understood, here we characterized the potential role of stomatin in this process. Stomatin is ubiquitously expressed in human tissue (Lapatsina et al., 2012). Targeting to the membrane is mediated by an *N*-terminal membrane insertion domain (Kadurin et al., 2009). Together with the stomatin-like protein 2 (SLP-2) stomatin forms hetero-oligomers at endosomal membranes (Mairhofer et al., 2009). In line, here by immunofluorescence we detected stomatin in the phagosomal membrane of RAW264.7 macrophages after infection with *A. fumigatus* conidia (Fig. 1). An identical observation was made with primary mouse macrophages (C57BL/6 BMDMs) (Fig. 1). These data indicate that conidia reside in phagosomes containing stomatin in their membrane.

**Figure 1:**
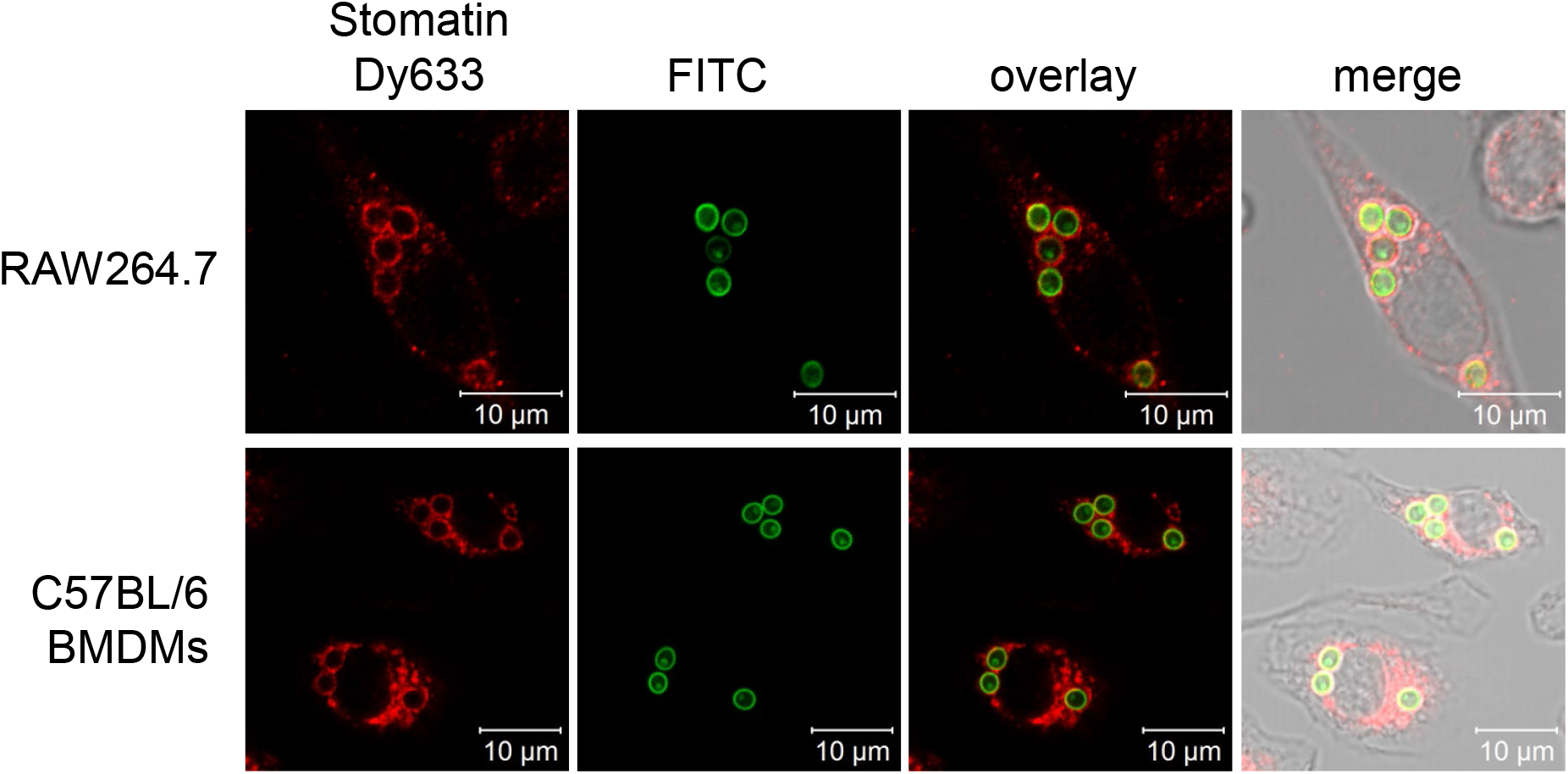
Localization of stomatin at phagosomal membranes of both RAW264.7 macrophages and bone marrow-derived macrophages (BMDMs) of C57BL/6 mice after infection with *A. fumigatus* conidia (FITC-labeled). Stomatin was detected by immunofluorescence using a stomatin antibody coupled to Dy633.

### CRISPR-Cas9 gene editing in RAW264.7 cells for generation of knockout cell lines

For functional analysis of stomatin, a knockout cell line for stomatin was created by CRISPR-Cas9 gene editing. First a stable macrophage cell line producing Cas9 was generated by transduction of RAW264.7 cells with Cas9 lentivirus particles. The expression of the Cas9 protein was confirmed by immunoblotting (Fig. 2A). These cells were further transduced with sgRNA lentivirus particles targeting exon 3 of the *Stom* gene. Hygromycin resistant clones were screened and the knockout of stomatin in selected clones was verified by Western blotting (Fig 2B). Sequence analysis of a selected clone that was used for all further studies, revealed deletion of an adenine at position 199 of the coding *Stom* sequence resulting in a frameshift and consequently in stomatin-deficient cells (Fig. 2C), as indicated by Western blot (Fig. 2B).

**Figure 2:**
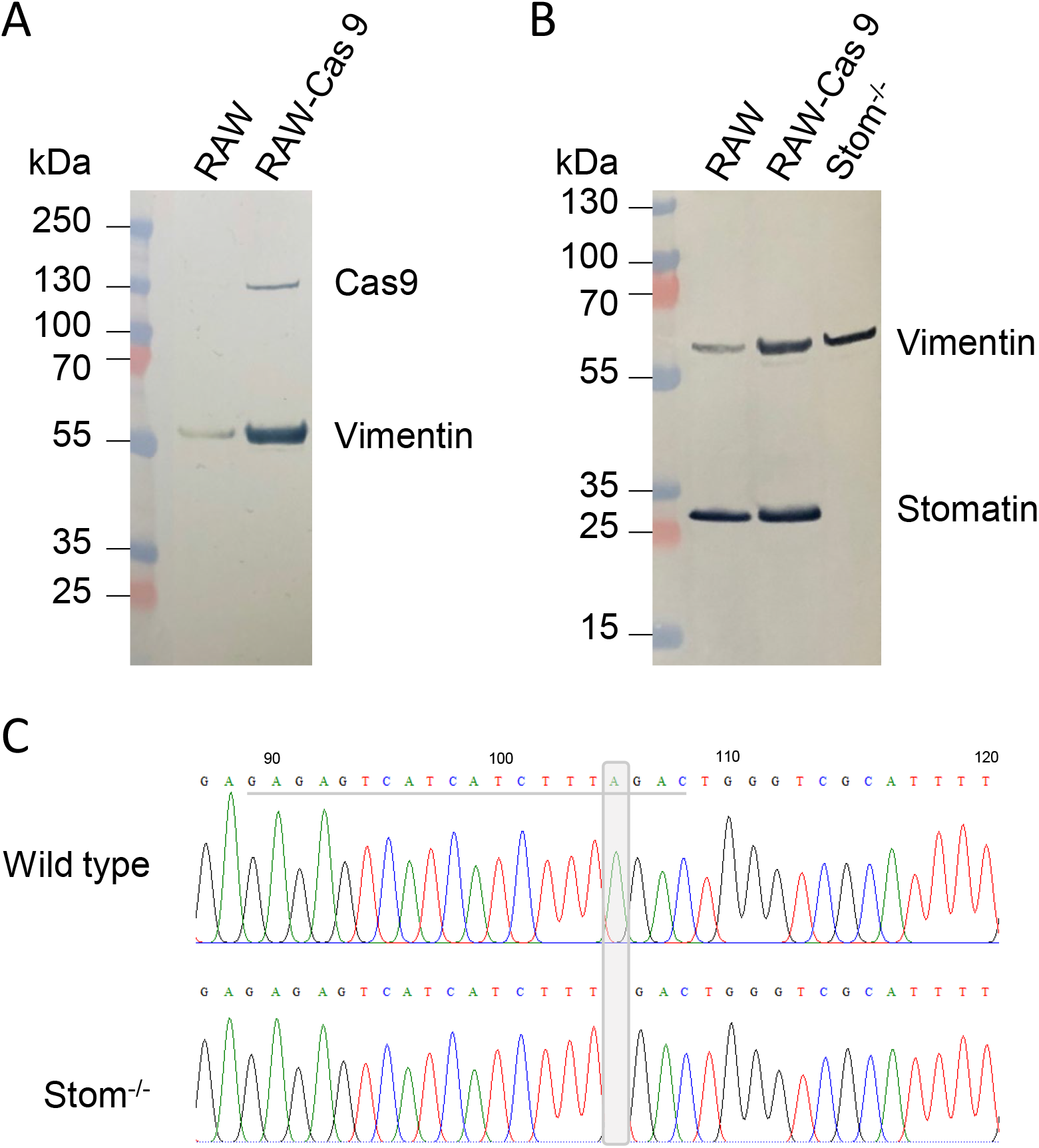
Generation of Stom^-/-^ cells. A) Identification of a RAW264.7 macrophage cell line with stable expression of Cas9. RAW264.7 cells were transduced with Cas9-lentivirus. A selected puromycin-resistant clone was analyzed by Western blot for production of Cas9. Vimentin was used as control. B) Western blot analysis to verify loss of stomatin protein in Stom^-/-^ RAW264.7 macrophages. C) Sequence analysis of Stom^-/-^ macrophages. CRISPR/Cas9 gene editing resulted in loss of an adenine (grey box) compared to the wild-type sequence. The sgRNA target sequence is underlined.

### Reduced phagocytosis of *A. fumigatus pksP* conidia in Stom^-/-^ macrophage cells

Phagocytosis depends on recognition of *A. fumigatus* by pattern recognition receptors and the reorganization of the membrane structure to form a phagocytic cup and to engulf the pathogen (Luther et al., 2008, Luther et al., 2007, Hasenberg et al., 2011). To analyze whether the knockout of stomatin affects phagocytosis, phagocytosis of FITC-labeled *A. fumigatus* conidia was monitored by immunofluorescence. As shown in Fig. 3A, in the Stom^-/-^ macrophage cells phagocytosis rate of *pksP* conidia was reduced by 50 % compared to wild-type cells. No effect on the phagocytosis rate of wild-type conidia was observed, since their DHN-melanin layer leads to reduced formation of functional lipid rafts and phagocytosis (Schmidt et al., 2020), Luther et al. 2007). Previously, it was shown that phagocytosis of *pksP* conidia depends on the pattern recognition receptor (PRR) dectin-1 that recognizes fungal beta-1,3-glucan (Luther et al., 2007, Steele et al., 2005). Therefore, it was reasonable to assume that stomatin is required for the recruitment of dectin-1 and we thus analyzed dectin-1 expression on membranes of macrophages by flow cytometry. However, we found no differences in the number of dectin-1-positive cells between wild-type and Stom^-/-^ macrophages (Figure 3B). Also, the median fluorescence intensity did not differ significantly between the two samples (Supplementary Figure 9). Besides recognition of fungal surface polysaccharides, dectin-1 is involved in phagosomal maturation processes (Mansour et al., 2013). Therefore, by immunofluorescence we determined the impact of stomatin on enrichment of dectin-1 on the phagosomal membrane surrounding conidia. Isolated phagosomes of RAW264.7 macrophages with internalized *pksP* conidia showed a strong dectin-1 signal (Fig. 3C). By contrast, phagosomes of Stom^-/-^ macrophages revealed a weak dectin-1 signal after phagocytosis of *pksP* conidia indicating that stomatin is required for quantitative accumulation of dectin-1 on the phagosomal membrane (Figure 3C).

**Figure 3:**
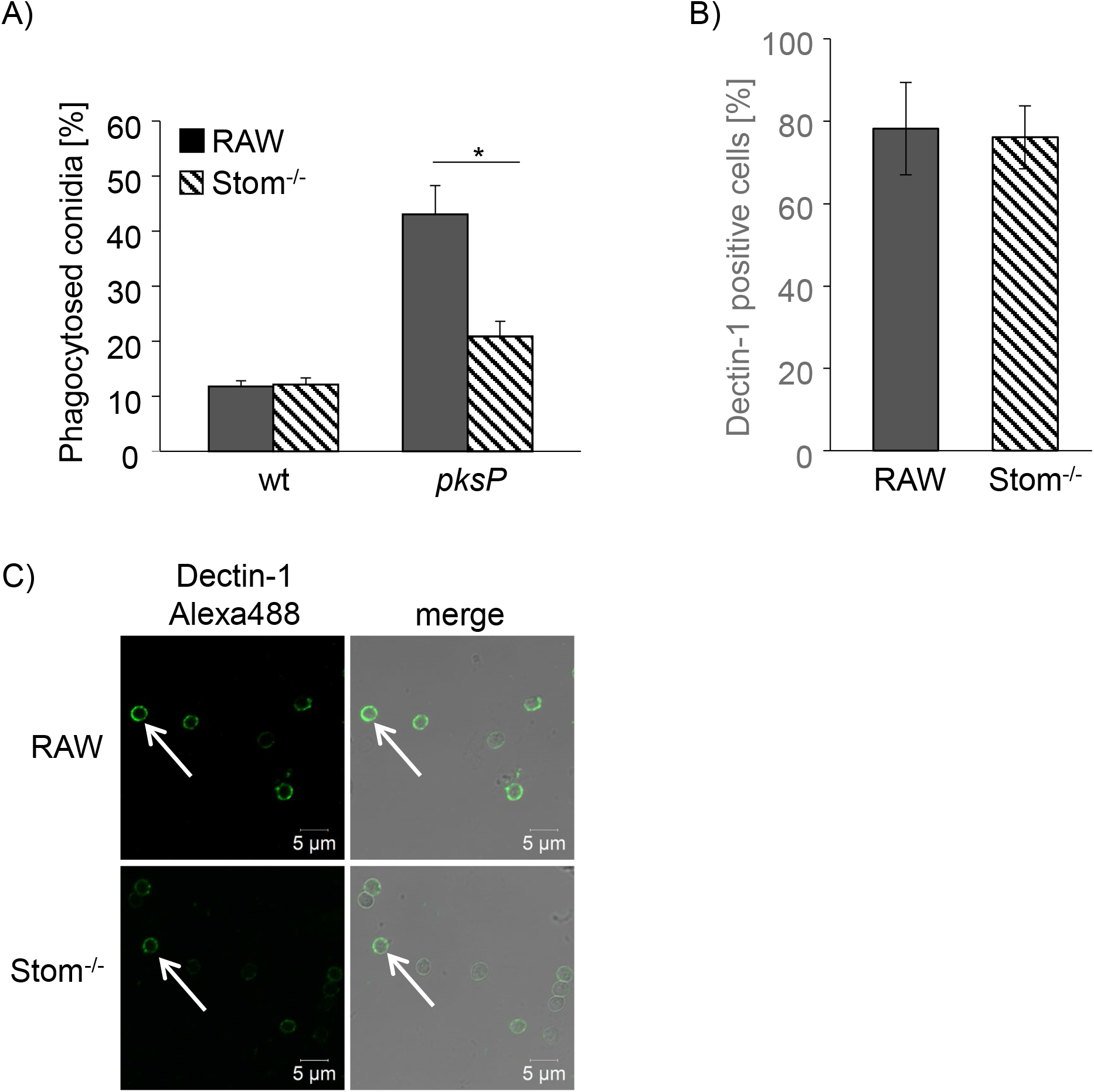
The role of stomatin in phagocytosis and accumulation of dectin-1. A) Percentage of phagocytosed *A. fumigatus* wild-type (wt) or *pksP* conidia by RAW264.7 and Stom^-/-^ macrophages. B) Percentage of dectin-1 positive cells measured by flow cytometry. Comparison of RAW264.7 wild-type and Stom^-/-^ macrophages. C) Immunofluorescence staining of dectin-1 on isolated phagosomes of wild-type and Stom^-/-^ macrophages after infection with *A. fumigatus pksP* conidia.

### Enrichment of sphingolipids in lipid-raft microdomains is independent of stomatin

To examine whether the enrichment of sphingolipids in lipid-raft microdomains is affected by stomatin, macrophages were infected with melanized wild-type and pigmentless *pksP*-mutant conidia and stained with Alexa Fluor 647-conjugated cholera toxin subunit B (CTB). CTB labels sphingolipid GM1 gangliosides. Accumulation of GM1 at distinct membrane sites is a characteristic feature of lipid rafts (Orlandi and Fishman, 1998). RAW264.7 wild-type as well as Stom^-/-^ macrophages that had engulfed wild-type conidia revealed only faint CTB staining of the phagosomal membrane. A stronger CTB signal was detected for *pksP* conidia-containing phagosomes in wild-type macrophages (Fig. 4A). By contrast, in Stom^-/-^ macrophages fainter CTB-mediated staining of the cytoplasmic membranes was detected. Counting of GM1-positive phagosomes revealed no significant differences between RAW264.7 wild-type or Stom^-/-^ macrophages (Fig. 4B).

**Figure 4:**
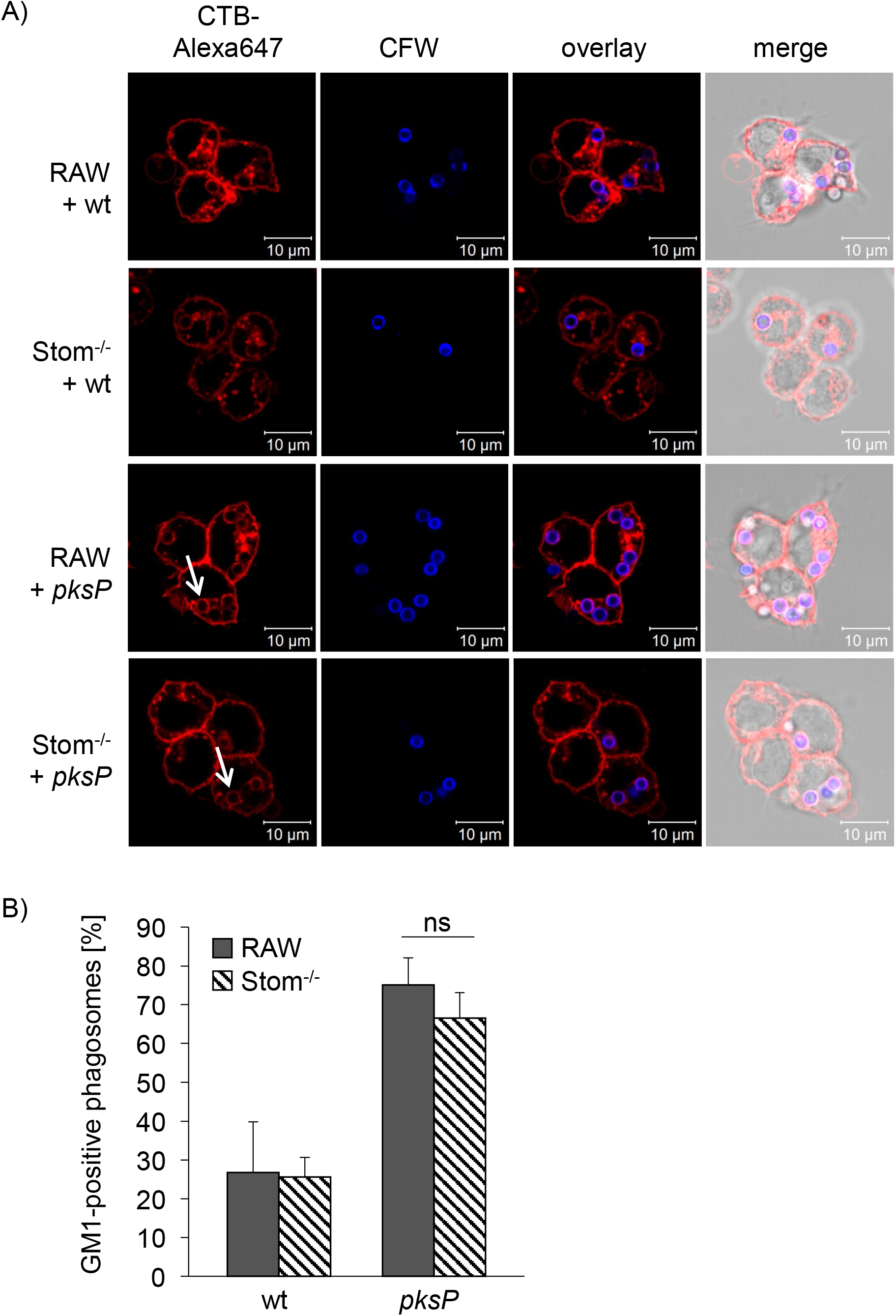
Enrichment of sphingolipids in lipid-raft microdomains. A) RAW264.7 and Stom^-/-^ macrophages were infected with CFW-labeled *A. fumigatus* wild-type or *pksP* conidia (CFW-labeled, blue) and stained for GM1 ganglioside (CTB-Alexa 647, red); white arrows indicate phagosomes. B) Percentage of GM1 positive phagosomes in RAW264.7 wild-type and Stom^-/-^ macrophages after infection with *A. fumigatus* wild-type or *pksP* conidia.

### Stomatin deficiency impairs vATPase assembly and interferes with acidification of conidia-containing phagosomes

It is known that in maturing phagosomes assembly of the vATPase complex takes place in membrane microdomains (Dermine et al., 2001, Lafourcade et al., 2008). This process depends on flotillin-dependent lipid rafts (Schmidt et al., 2020). To determine whether stomatin is also involved in vATPase assembly, the percentage of the assembled protein complexes at the phagosomal membranes was quantified by immunofluorescence. As shown in Figure 5A, the fluorescence due to the amount of subunit V_1_ assembled to the membranous V_0_ complex on the phagosomal membrane was affected by stomatin (Fig. 5A). In comparison to *pksP*-containing phagosomes of RAW264.7 macrophages, which display homogenous vATPase assembly at the phagosomal membrane (Schmidt et al., 2020), the distribution of the signal at phagosomal membranes of Stom^-/-^ cells appeared more fragmented. Phagosomes that contained wild-type conidia only showed faint staining of the phagosomal membrane in comparison to *pksP* conidia surrounding phagosomes in both cell lines. Image-based analyses of the signal intensities verified these findings. Phagosomes of Stom^-/-^ macrophages showed reduced fluorescence intensities after infection with *pksP* conidia compared to phagosomes from RAW264.7 wild-type cells (Fig. 5B), indicating a reduced number of vATPase in the phagosomal membrane of Stom^-/-^ macrophages.

**Figure 5:**
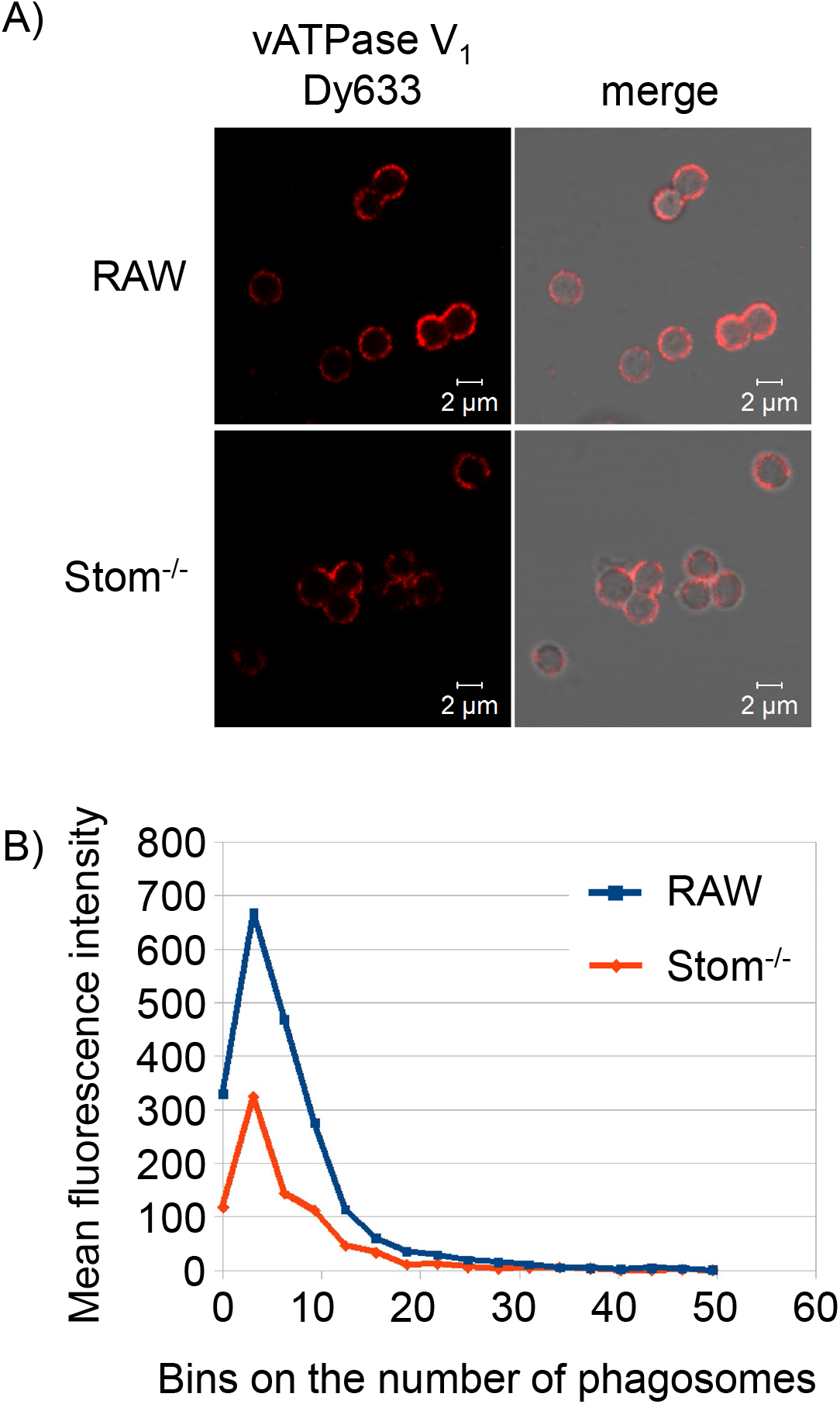
Impact of stomatin on vATPase assembly and acidification of phagosomes. A) vATPase V_1_ assembly in RAW264.7 and Stom^-/-^ macrophages after infection with *A. fumigatus pksP* conidia detected by immunofluorescence of V_1_ subunit. B) Quantification of the mean fluorescence intensities for the vATPase of isolated phagosomes containing *pksP* mutant conidia.

vATPase-dependent acidification of phagosomes can be detected by using LysoTracker Red (Thywissen et al., 2011) that shows red fluorescence of phagosomes. Here, a difference in acidification of phagosomes between wild-type and Stom^-/-^ macrophages was observed (Fig. 6A, B). As previously reported (Thywissen et al., 2011), virtually no fluorescence of phagosomes containing wild-type conidia was observed, in contrast to clear fluorescence seen in phagosomes containing *pksP* conidia. For stomatin-deficient macrophages, fluorescence of some *pksP* conidia-containing phagosomes was reduced (Fig. 6A, arrows). To verify these data, we employed an additional sensor dye, *i.e*., LysoSensor Yellow/Blue, which allows for visualization of gradual changes in pH in phagosomes. Depending on the pH, its emission wavelength shifts from blue (pH neutral) to yellow (acidic). RAW264.7 macrophages infected with *pksP*-conidia showed bright yellow rings at the phagosome indicating full acidification (Fig. 6C). By contrast, in Stom^-/-^ cells only some phagosomes stained yellow, but others remained blue or displayed a dotted ring-like structure, indicating impaired acidification.

**Figure 6:**
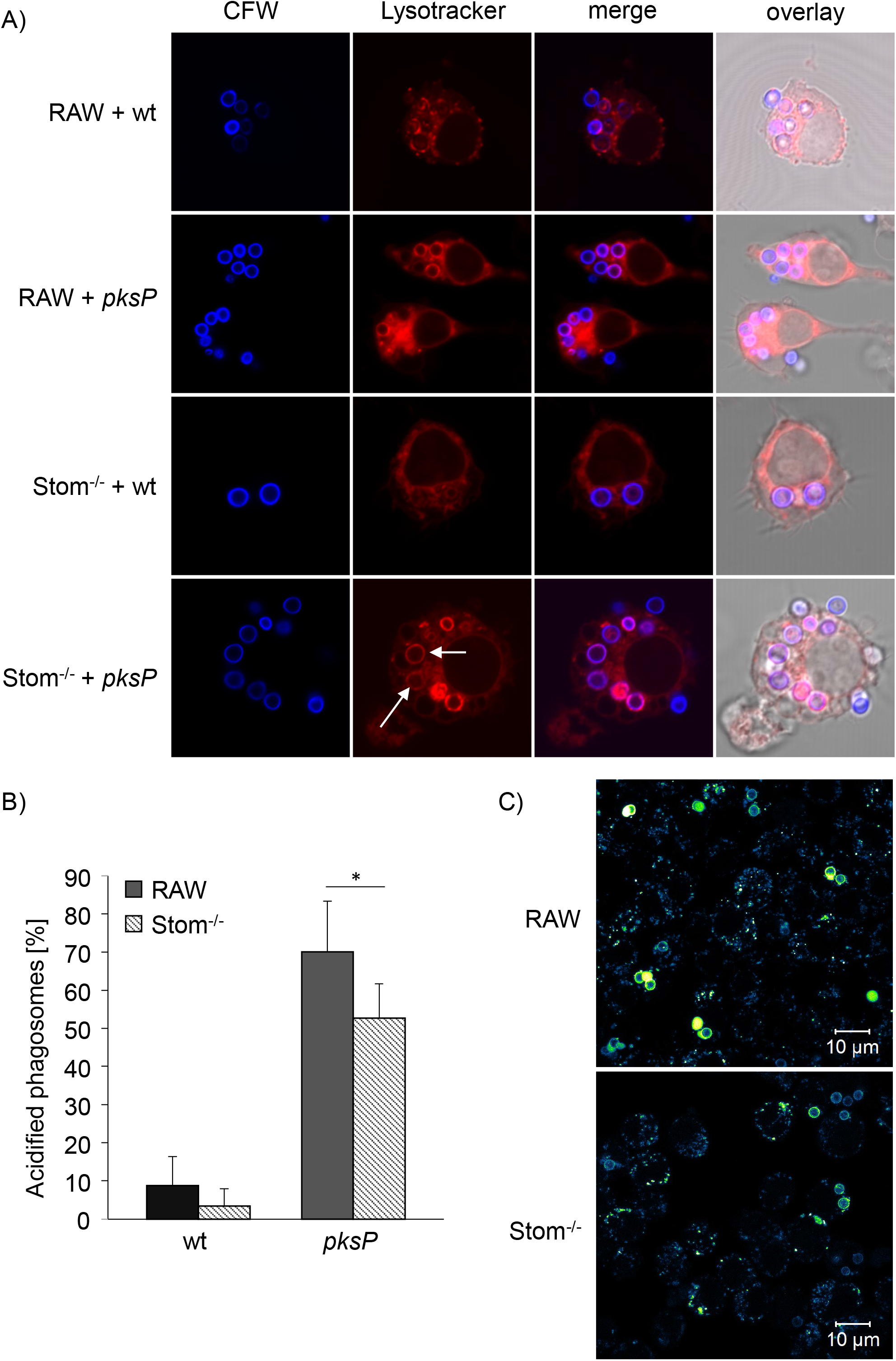
Impairment of phagosomal acidification by stomatin deficiency. A) Detection of acidified phagosomes of RAW macrophages containing wild-type (wt) or *pksP* conidia using LysoTracker Red. Conidia were stained with CFW (blue). B) Quantification of A. C) Staining of phagosomes containing *pksP* conidia (stained with CFW, blue) in Stom^-/-^ cells with LysoSensor.

### Stomatin is required for maturation of phagosomes to phagolysosomes

Due to the impaired acidification and reduced vATPase assembly at the phagosomal membrane we hypothesized an involvement of stomatin in the maturation process of phagosomes. However, acidification of phagosomes in the stomatin knockout was less impaired when compared to that seen in a double knockout of flotillins 1 and 2 (Schmidt et al., 2020). Therefore, we monitored additional phagosomal maturation parameters such as the fusion of lysosomes with phagosomes. Such an analysis is feasible by employing tetramethylrhodamine dextran-loaded lysosomes. A successful fusion of lysosomes with phagosomes is indicated by a rhodamine signal appearing in the phagosome. As shown in Figure 7A and B, a significantly reduced rhodamine signal in phagosomes was observed for Stom^-/-^ macrophages compared to wild-type RAW264.7 cells after infection with *pksP* mutant conidia. As expected, because of the disruptive effect of DHN-melanin on the formation of lipid-raft microdomains and thus maturation of phagosomes, infection with melanized wild-type conidia resulted in a low rhodamine signal in phagosomes in both, RAW264.7 wild-type cells and Stom^-/-^ macrophages.

**Figure 7:**
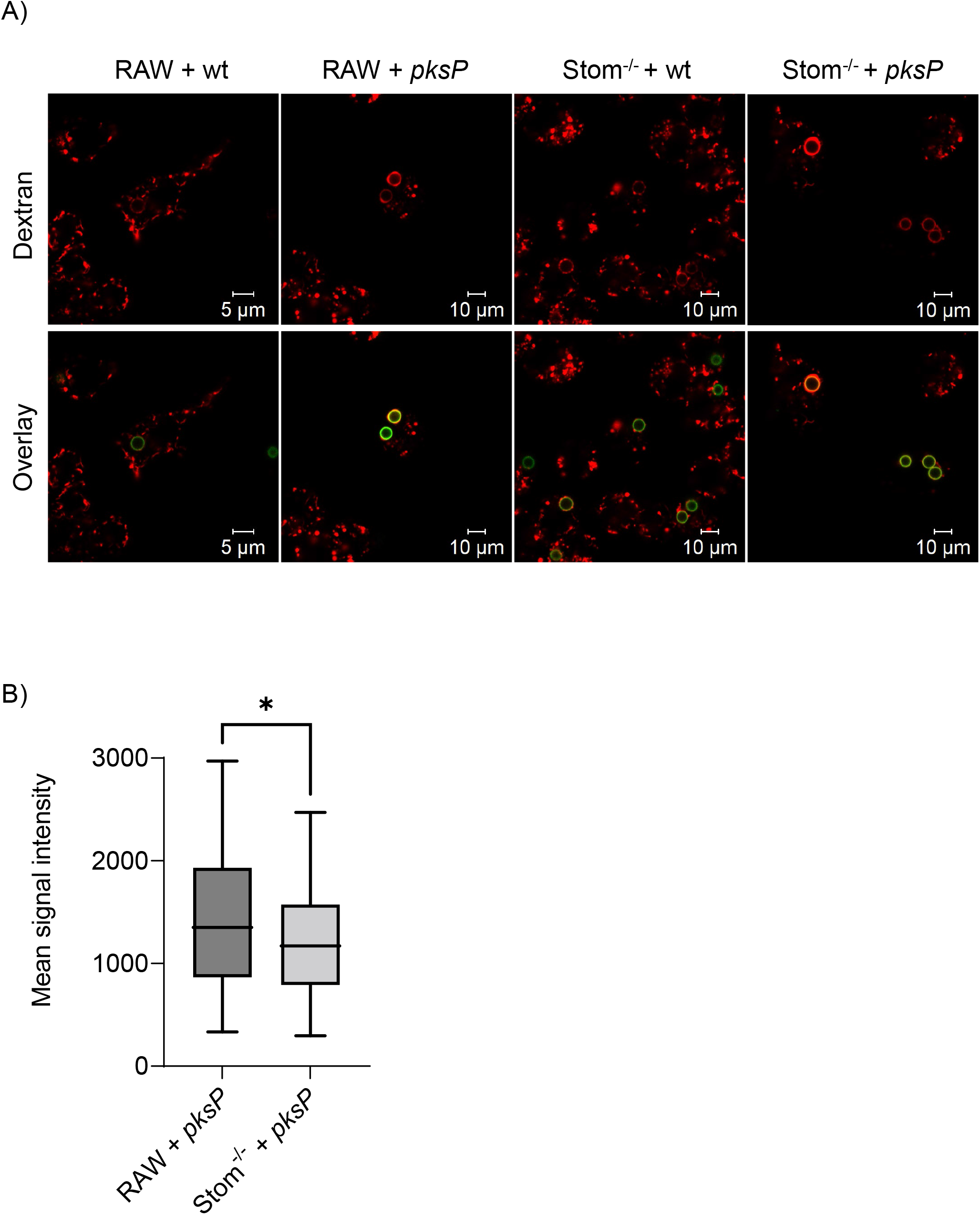
Evaluation of phagosomal maturation. A) Macrophages infected with FITC-labeled *pksP* conidia. The tetrarhodamine dextran stain at the phagosome indicates fusion with lysosomes and maturation to phagolysosomes. B) Comparison of mean fluorescence intensities of *pksP* conidia-containing phagosomes in RAW264.7 or Stom^-/-^ cells.

## Discussion

Lipid rafts are small, highly dynamic microdomains in membranes that recruit and concentrate molecules involved in cellular signaling. In accordance with this function, membrane microdomains are also required for antifungal immunity (Varshney et al., 2016, Schmidt et al., 2019, Schmidt et al., 2020). A prominent and ubiquitously found protein in lipid-raft microdomains is stomatin that was suggested to be involved in the reorganization of the cytoplasmic membrane structure (Salzer and Prohaska, 2001, Snyers et al., 1999, Snyers et al., 1998). Stomatin is known to interact with various ion channels and transporters thereby modulating their activities (Unfried et al., 1995, Umlauf et al., 2006, Snyers et al., 1999, Brand et al., 2012, Gallagher and Forget, 1995). Here, our data fully agree with the localization of stomatin in various cellular membranes. Immunofluorescence analyses revealed accumulation of stomatin on both the cytoplasmic and phagolysosomal membrane of both primary macrophages derived from bone marrow and RAW264.7 macrophages. This data is also in line with proteomic studies of isolated phagolysosomes from RAW264.7 macrophages identifying stomatin (Schmidt et al., 2018).

We also discovered that stomatin is important for the phagocytosis of *A. fumigatus* conidia. This conclusion was drawn of experiments based on the successful inactivation of the stomatin gene in the macrophage cell line RAW264.7. In Stom^-/-^ macrophages, reduced phagocytosis was detected for *pksP* conidia. For wild-type conidia, there was no difference in phagocytosis between Stom^-/-^ and wild-type cells. This observation agrees well with previous results that DHN-melanin of wild-type conidia covers immunogenic structures and are therefore less accessible for receptors in wild-type conidia (Heinekamp et al., 2015). Thus, an effect by lack of the lipid-raft microdomain component stomatin in association with melanized wild-type conidia was not expected.

An important receptor recognizing conidia is the C-type lectin receptor dectin-1 that recognizes the fungal surface polysaccharide β-1,3-glucan, which is more accessible on *pksP* conidia than on melanized wild-type conidia (Meier et al., 2003, Luther et al., 2007, Chai et al., 2011). Presence of dectin-1 increases phagocytosis of *pksP* conidia and induces the production of cytokines and chemokines triggering an inflammatory response (Chai et al., 2010).

Dectin-1 was also shown to be a regulator of phagosomal maturation (Mansour et al., 2013). Our data support this conclusion, since we found that stomatin also contributes to the enrichment of dectin-1 on the phagosomal membrane after phagocytosis of *A. fumigatus* conidia. Furthermore, as shown here, stomatin is required for full acidification of phagolysosomes. Acidification of the phagolysosomal lumen depends on the vATPase proton pump (Cotter et al., 2015). A direct link between lipid-raft microdomains and assembly of the vATPase on the phagosomal membrane has been already reported (Dhungana et al., 2009, Lafourcade et al., 2008, Schmidt et al., 2019, Schmidt et al., 2020). The vATPase complex was shown to co-localize with the lipid-raft marker protein flotillin-1 (Ryu et al., 2010, Schmidt et al., 2019, Schmidt et al., 2020). Therefore, it seemed likely that stomatin is also required for the assembly and or enrichment of the vATPase complex in maturing phagosomes. This assumption was proven by our immunofluorescence studies revealing that stomatin is involved in the recruitment of the V_1_ subunit to the membranous V_0_ subunit to form a functional vATPase complex and consequently, functional vATPase can pump protons into the phagosomal lumen. We speculated whether the assembly of vATPase was disturbed when stomatin was lacking or rather fusion of phagosomes and lysosomes. Our data on measuring the fusion events are in favor of reduced fusion of phagosomes and lysosomes.

Previously, we reported that conidial DHN-melanin interferes with formation of flotillin-dependent lipid-raft microdomains in the phagolysosomal membrane (Schmidt et al., 2020). Surprisingly, the localization of stomatin on the membrane was not affected by DHN-melanin and also lack of stomatin did apparently not lead to major changes in lipid rafts because staining for GM1 in Stom^-/-^ cells did not differ from that seen in wild-type cells. Therefore, an obvious question concerns the link between stomatin and flotillins. Both lipid raft markers belong to the SPFH domain proteins (Browman et al., 2007) and share various characteristics like related topology and regulatory roles for signaling pathways (Schulte et al., 1997, Bickel et al., 1997, Arkhipova et al., 2014, Baumann et al., 2000). Like stomatin, flotillins are ubiquitously expressed and are conserved proteins involved in several cellular processes as membrane trafficking, phagocytosis, phagosomal maturation and T cell activation (Babuke and Tikkanen, 2007). Our data suggest that stomatin does not function as chaperon like flotillins or caveolin that apparently structure lipid-raft microdomains but rather is required for positioning of certain proteins like dectin-1 in lipid rafts. In line with our hypothesis, in lipid-raft microdomains we observed co-localization of stomatin and flotillin at the phagolysosomal membrane of RAW264.7 macrophages after phagocytosis of *A. fumigatus* conidia (Supplementary Figure 10).

Collectively, our data are in accordance with the following model: In stomatin knockout cells, lipid-raft microdomains are still present. Stomatin is likely important for positioning of certain proteins like dectin-1 in lipid raft. It is likely that this is achieved by a stomatin-dependent production of larger lipid-raft microdomains, that, in addition, allow for more efficient fusion of phagosomes with lysosomes and thus for an increase of vATPase and dectin-1 molecules on the mature phagolysosomes. In line, accumulation of dectin-1 on phagosomal membranes was shown to be dependent of acidification of phagosomes (Mansour et al. 2013) and dectin-1 co-localized to lipid-raft microdomains (Xu et al., 2009). Alternatively, stomatin could be involved in the direct fixation of proteins in lipid raft microdomains that is in part triggered by the fusion of phagosomes with lysosomes.

## Conclusion

The data presented here provide new insights on the important role of the integral membrane protein stomatin in the immune response against human pathogenic fungi. We provide evidence that stomatin plays a crucial role for the quantitative localization of vATPase and dectin-1 in both cytoplasmic and phagosomal membranes, thereby impacting phagocytosis and intracellular processing of *A. fumigatus* conidia.

## Acknowledgments

This work was funded by the Deutsche Forschungsgemeinschaft (DFG, German Research Foundation) through the Collaborative Research Center/Transregio 124 “FungiNet”, DFG project number 210879364 (projects A1, and B4) and the SFB 1278 “PolyTarget”, DFG project number 316213987 (project Z01), and the Cluster of Excellence 2051 “Balance of the Microverse”, DFG project number 390713860. FS and MG were members of the excellence graduate school Jena School for Microbial Communication (JSMC) funded by the DFG and the International Leibniz Research School for Microbial and Biomolecular Interactions (ILRS) as part of the JSMC, respectively. TO was paid by the Austrian Science Fund (FWF) through the Erwin Schrödinger Fellowship (project number J4432).

## Data availability statement

All data and materials that support the results or analyses presented in this paper are freely available under DOI:10.6084/m9.figshare.19122008.

## Supplementary Figure Legends

**Supplementary Figure 1.** The compartment structure of the phagolysosome analysis workflow in JIPipe. The left column shows the two compartments responsible for training the default Cellpose model further, in order to improve its predicted segmentation performance. The top compartment prepares the annotated images for Cellpose transfer learning, whereas the bottom compartment executes the actual training (details are shown in Supplementary Figure 2.) In the right column the two compartments are shown that apply both the default and transfer learning—based Cellpose models to segment the phagolysosome images (top panel) and carry out postprocessing (e.g., watershed algorithm to resolve phagolysosome clusters) if necessary (bottom panel).

**Supplementary Figure 2.** The workflow of the Train Cellpose JIPipe compartment, which trains the default Cellpose model further by using transfer learning. Images were read into memory and the folder and subfolder names, as well as replicate and strain identifiers were saved as JIPipe annotations (black round-corner rectangle). Images manually annotated with regions of interest (ROIs) designating phagosomes that were not recognized by the default Cellpose model were read from selected folders (black triangle). The annotated JIPipe objects were separated into images and regions of interest (black rectangles), cut into 256×256 pixel tiles (black rhomboid), which were fed into a JIPipe node that was designed to provide a graphical way to train the Cellpose model via transfer learning (green oblique rectangle). The trained model (blue rectangle) was saved and used in the compartment designed for applying default or trained Cellpose models for phagolysosome segmentation (blue oval; see Supplementary Figure 3).

**Supplementary Figure 3.** The JIPipe workflow of applying the Cellpose model to segment phagosomes in transmitted light images and to measure the corresponding vATPase fluorescence in these regions of interest. The image properties (folders and subfolders, replicate and strain names, as well as the image filename) were identified and added to the image objects in the form of JIPipe annotations (black round-corner rectangle). The Scale images step (black rhomboid) unified the pixel size in all images, a step made necessary by having confocal images of various spatial resolutions. The Channel assignment step (black rhomboid) separates the transmitted light (grey rectangle) and fluorescence channels (red rectangles), the latter corresponding to the vATPase signal. The transmitted light images were then corrected for illumination inhomogeneities (grey rhomboid) and plugged into the Cellpose application nodes (green oblique rectangles). Here one Cellpose node each was used to apply the default model and the transferred learning-based model, respectively. The latter received the trained model from the Cellpose training compartment (blue oval and rectangle) as shown in Supplementary Figure 2. The segmented phagolysosomes were filtered based on their size and shape (grey hexagons): only objects between the size of 70 and 500 pixels, and with circularity above 0.5 were considered for further analysis. Since each image had two segmentations (default and transfer learning model—based), a manual decision step was inserted as a next step (grey diamond), to decide which model to use for the measurements. The selected model provided the ROIs (magenta wavy rectangle) that were used in the next step to measure the vATPase fluorescence intensity in the phagolysosomes (black oval). The intensity values within the ROIs projected onto the vATPase image (red rhomboid) were saved for further analysis, including intensity distribution histograms under various conditions.

**Supplementary Figure 4.** Illustrative images demonstrating the selection between the default and transfer learning—based trained Cellpose models. In the horizontal pairs on images, the left and right panels show the segmentation predicted by the default or transfer learning— based Cellpose models, respectively (yellow outlines). In (A) and (B) the default model outperformed the trained model, whereas in (C)-(D) and (E)-(F) transfer-learning was necessary to recognize the phagolysosomes that appear in a darker phase in the transmitted light images.

**Supplementary Figure 5.** The node array of the Cellpose preprocessing compartment in Supplementary Figure 1 left, top panel. The functional steps of the node arrays in Supplementary Figure 1 left are detailed in Supplementary Figure 2.

**Supplementary Figure 6.** The node array of the Cellpose training compartment in Supplementary Figure 1 left, bottom panel. The functional steps of the node arrays in Supplementary Figure 1 left are detailed in Supplementary Figure 2.

**Supplementary Figure 7.** The node array of the Apply Cellpose compartment in Supplementary Figure 1 right, top panel. The functional steps of the node arrays in Supplementary Figure 1 right, top panel, are detailed in Supplementary Figure 3.

**Supplementary Figure 8.** The detailed node view of the manual decision step in Supplementary Figure 3 (grey diamond).

**Supplementary Figure 9.** Median Fluorescence intensity of dectin-1 of RAW264.7 macrophages and Stom^-/-^ cells measured by flow cytometry.

**Supplementary Figure 10.** lmmunofluorescence to monitor co-localization of flotillin-1 and stomatin on isolated phagolysosomes of macrophages after infection with *pksP* conidia.

