## Supplementary figures and images for "The lipid raft-associated protein stomatin is required for accumulation of dectin-1 in the phagosomal membrane and for full activity of macrophages against *Aspergillus fumigatus*"

### Supplemental Figures

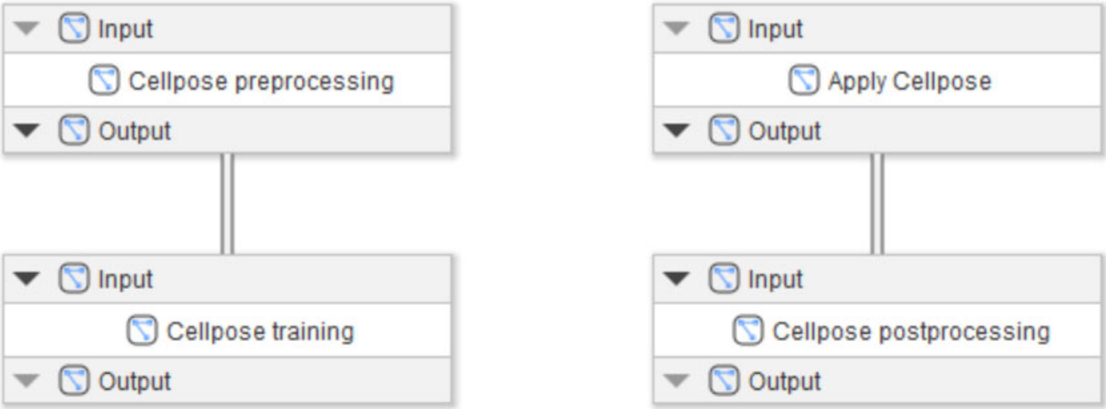

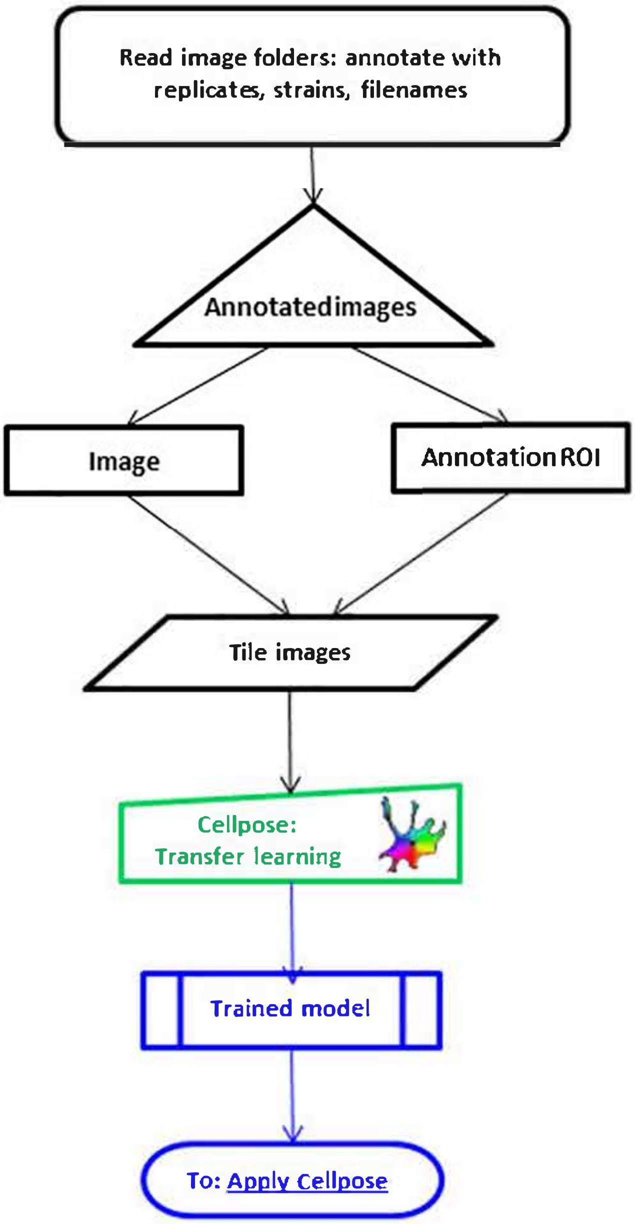

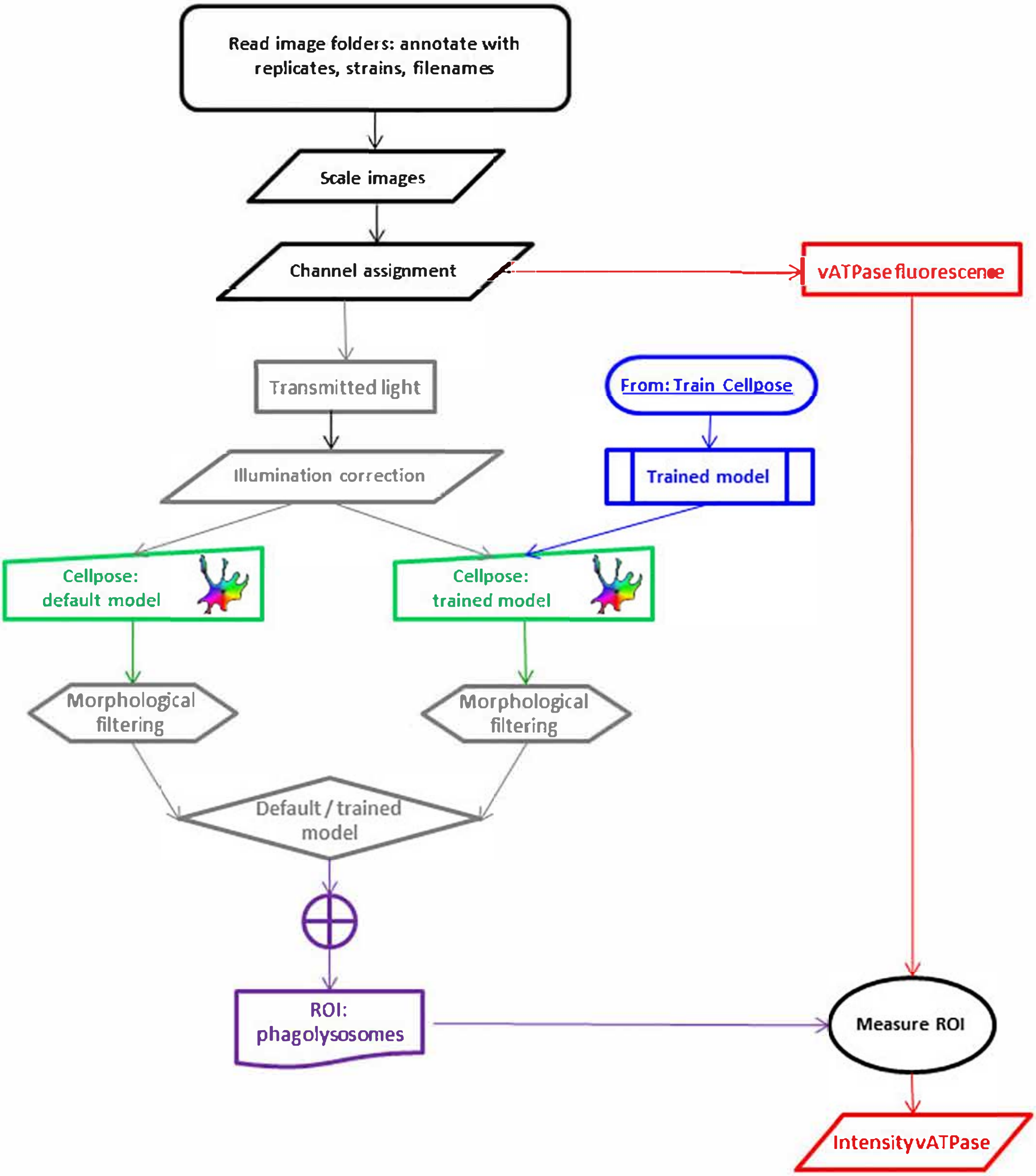

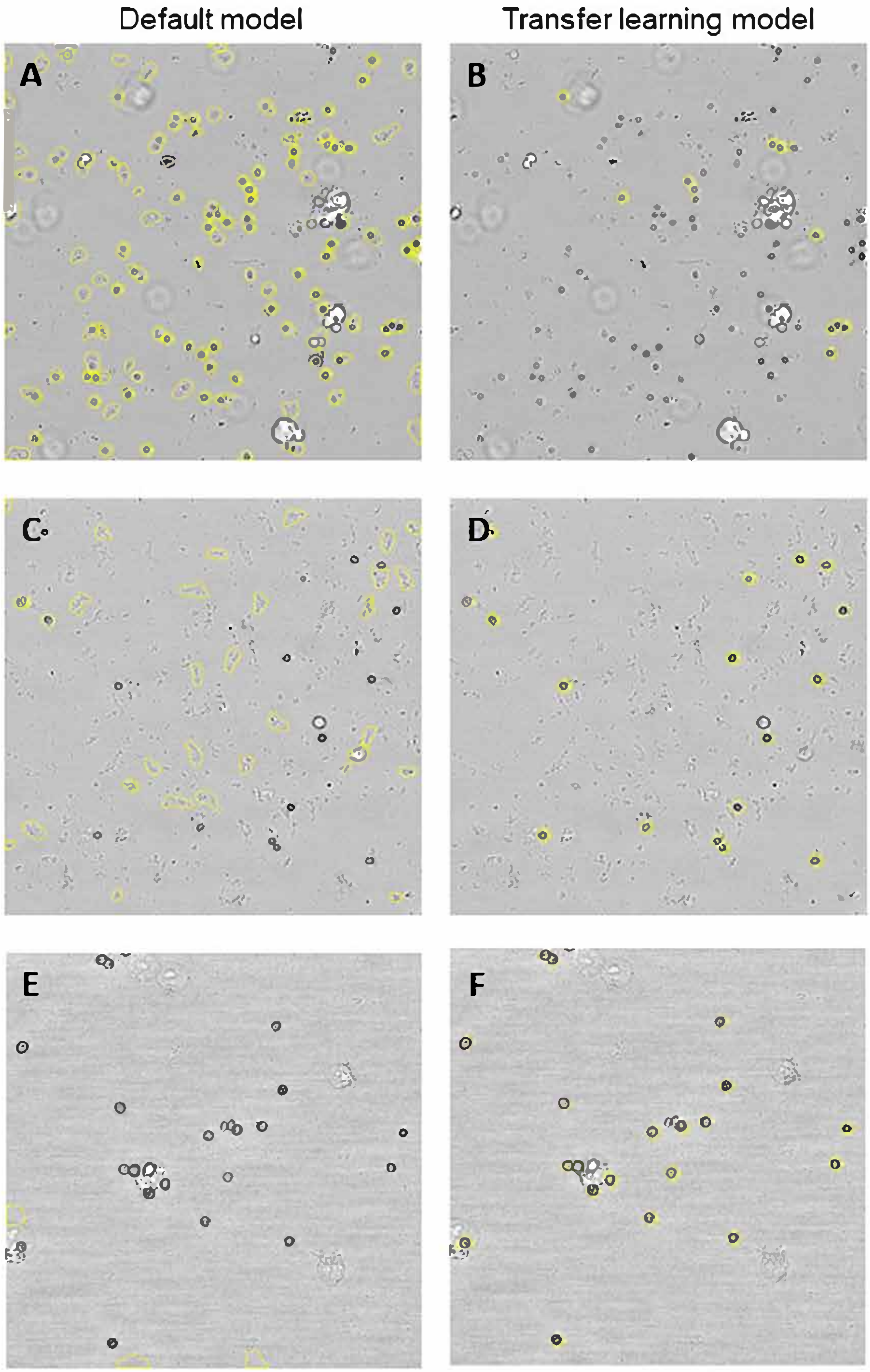

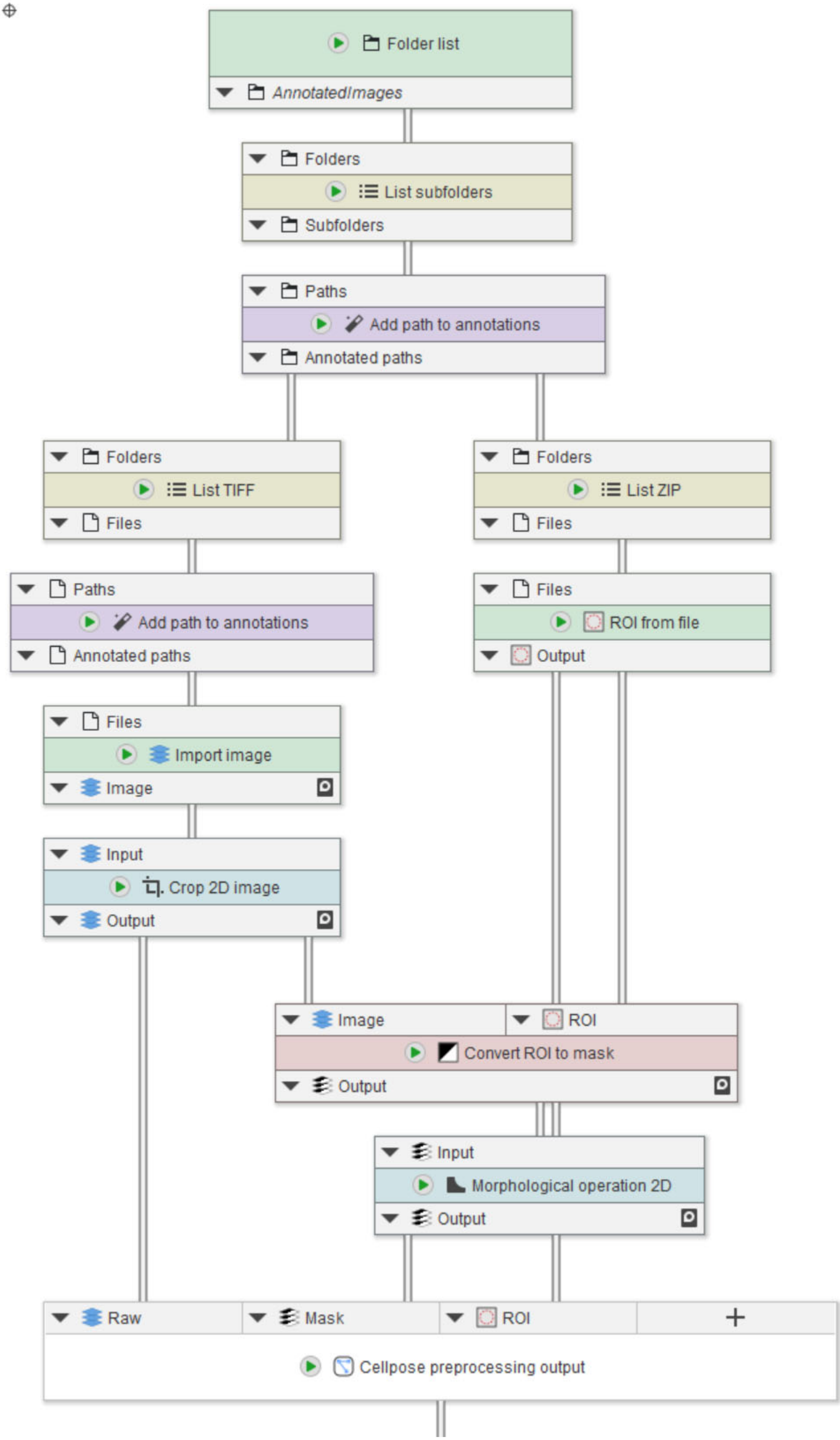

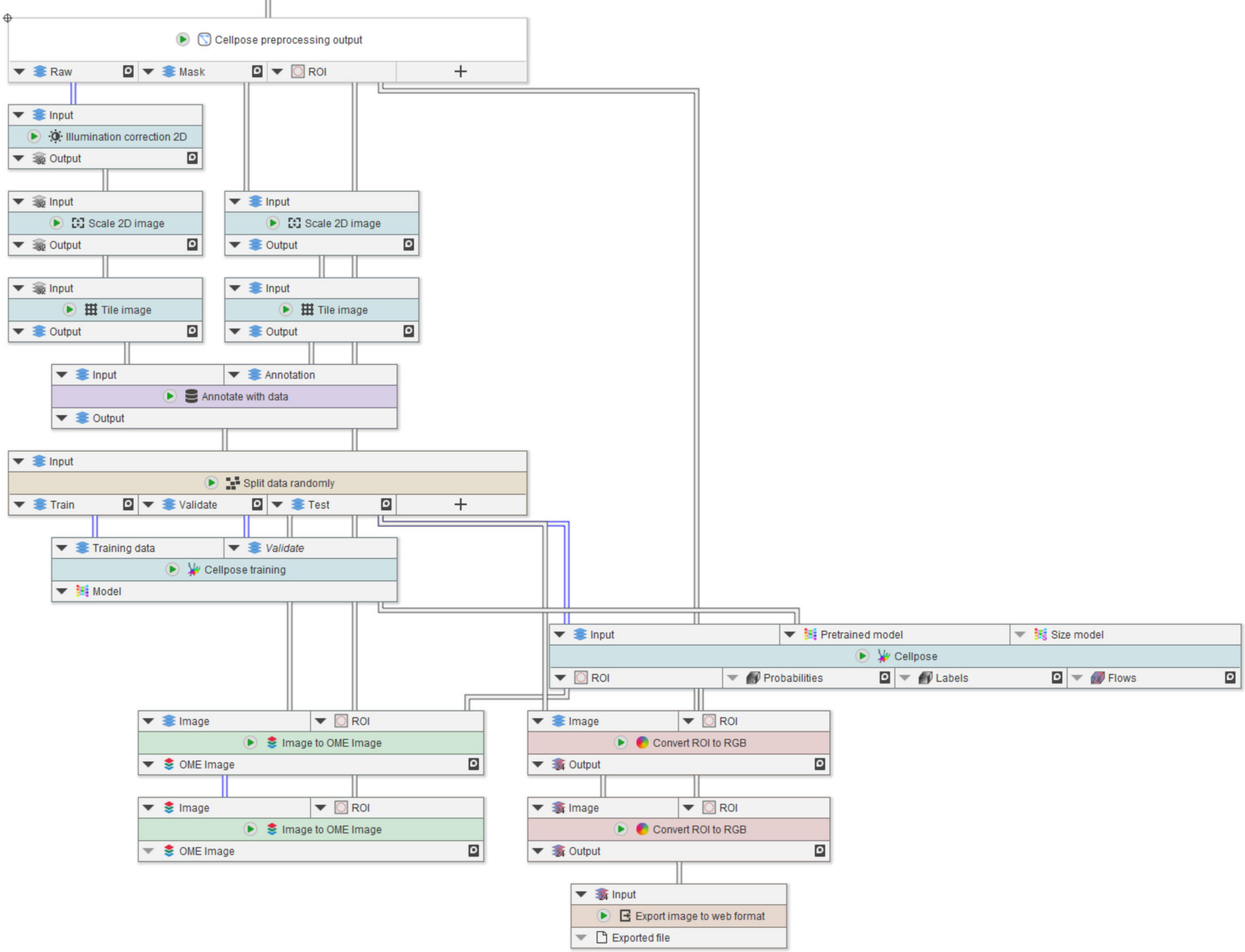

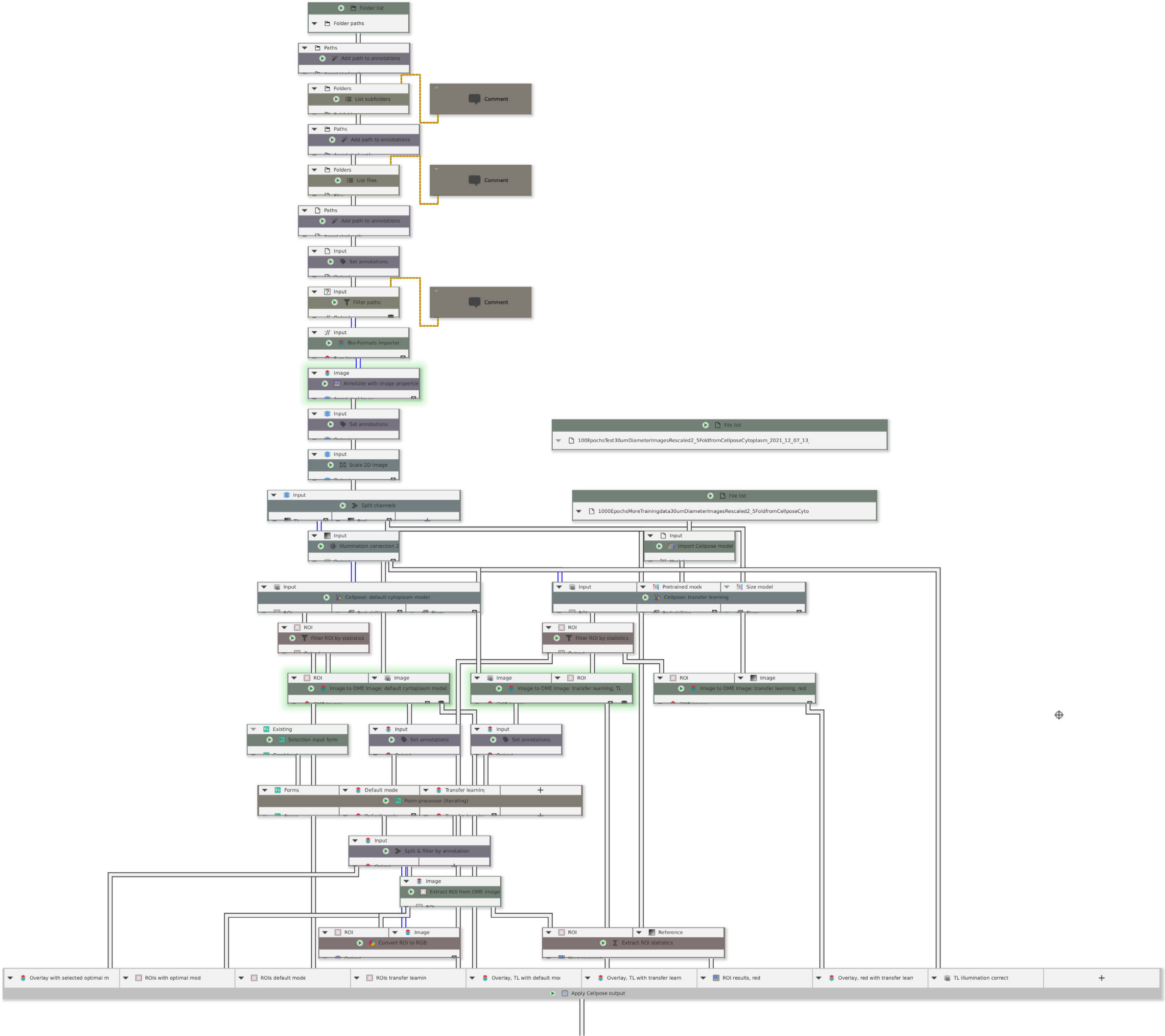

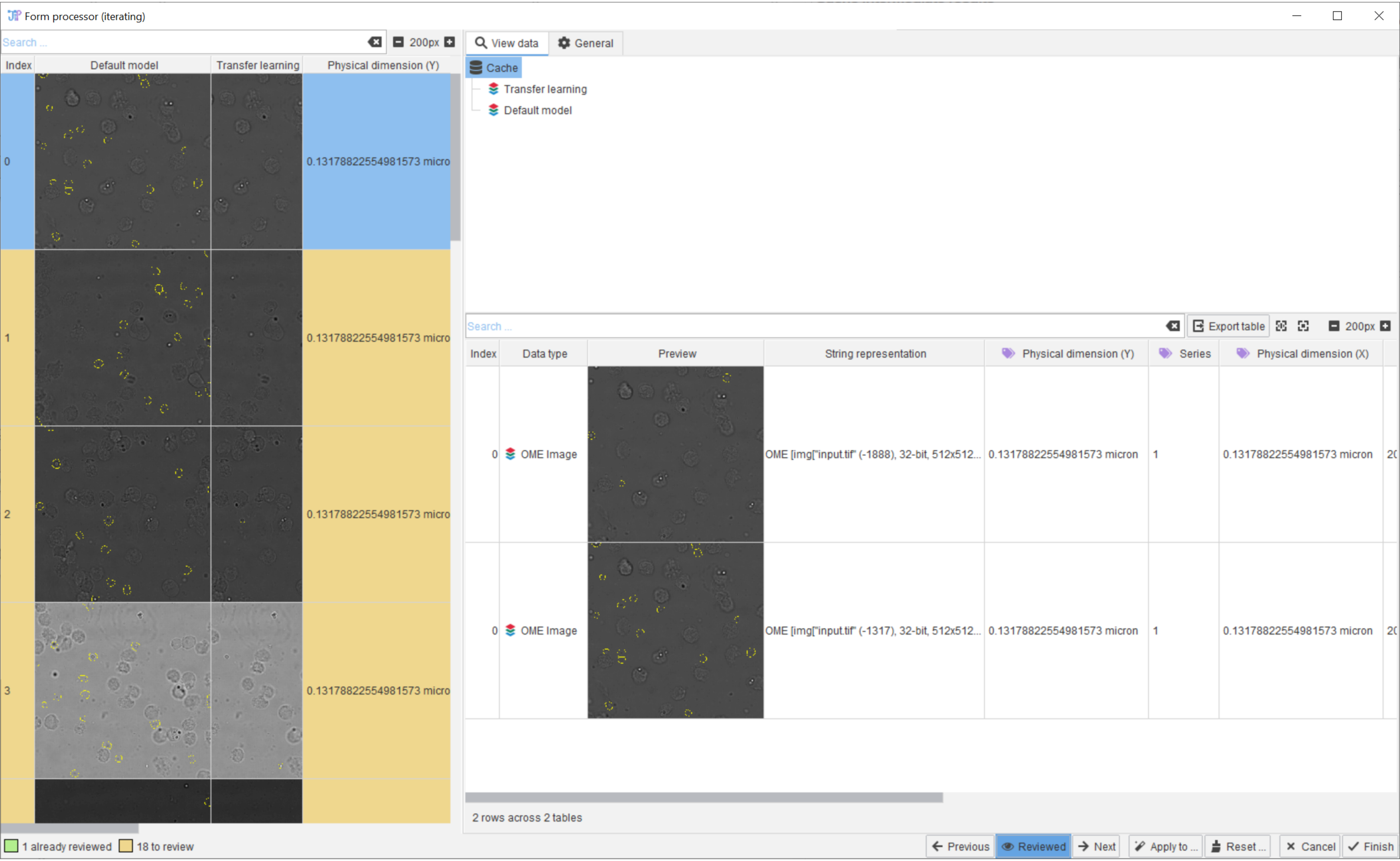

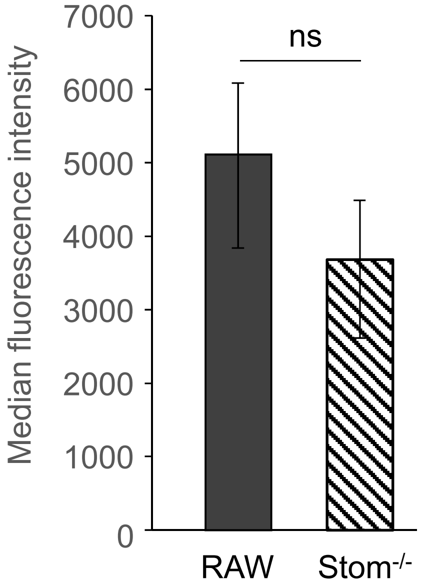

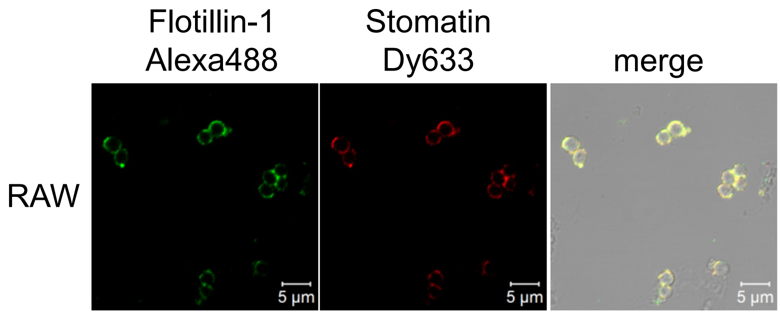
